# Control of synaptic specificity by limiting promiscuous synapse formation

**DOI:** 10.1101/415695

**Authors:** Chundi Xu, Emma Theisen, Elijah Rumbaut, Bryan Shum, Jing Peng, Dorota Tarnogorska, Jolanta A. Borycz, Liming Tan, Maximilien Courgeon, Ian A. Meinertzhagen, Matthew Y. Pecot

## Abstract

The ability of neurons to distinguish appropriate from inappropriate synaptic partners in their local environment is fundamental to the proper assembly and function of neural circuits. How synaptic partner selection is regulated is a longstanding question in Neurobiology. A prevailing hypothesis is that appropriate partners express complementary molecules that match them together and promote synaptogenesis. Dpr and DIP IgSF proteins bind heterophilically and are expressed in a complementary manner between synaptic partners in the *Drosophila* visual system. Here, we show that in the lamina, DIP mis-expression is sufficient to promote synapse formation with Dpr-expressing neurons, and that DIP proteins are not necessary for synaptogenesis but rather function to prevent ectopic synapse formation. These findings indicate that Dpr-DIP interactions regulate synaptic specificity by biasing synapse formation towards specific cell-types. We propose that synaptogenesis occurs independent of synaptic partner choice, and that precise synaptic connectivity is established by limiting promiscuous synapse formation.

## INTRODUCTION

The nervous system comprises tremendous cellular diversity, yet the ability of organisms to interact appropriately with their environment depends on neurons establishing precise patterns of synaptic connections. How organisms navigate cellular complexity and correctly assemble their nervous systems with high fidelity and precision is a fundamental question in Neurobiology. In general, precise neural connectivity is thought to be established in steps. For example, targeting events such as axon guidance, topographic positioning, and laminar innervation progressively restrict the partners available for synapse formation. However, in their local environment neurons still face the challenge of identifying correct synaptic partners amidst many alternatives, referred to here as synaptic specificity. Based on their landmark experiments showing that regenerating neurons have the capacity to select appropriate targets in the face of many choices, Langley (Langley, 1895) and Sperry (Sperry, 1963) proposed that there must be a specific molecular relationship between appropriate synaptic partners that allows them to identify each other and form synapses in a complex environment. A common interpretation of this idea is that synaptic partners express complementary cell recognition molecules that match them together through a lock and key-like mechanism and promote synaptogenesis. In the past decades numerous cell surface molecules have been found to regulate synaptic connectivity (Missler et al., 2012; Sudhof, 2017; Yamagata et al., 2003). However, these predominantly regulate the patterning of axons and dendrites, synapse structure and function, and few are known to control synaptic specificity.

Recent biochemical and gene expression studies have demonstrated that the members of two subfamilies of the Immunoglobulin superfamily (IgSF), the Dpr (defective proboscis retraction) family (21 members) (Carrillo et al., 2015; Nakamura et al., 2002) and the family of dpr-interacting proteins (DIPs) (9 members) (Ozkan et al., 2013), bind heterophilically (Carrillo et al., 2015; Ozkan et al., 2013) and are expressed in a complementary manner between synaptically coupled cell types in the *Drosophila* visual system (Carrillo et al., 2015; Tan et al., 2015). Based on these findings, it was proposed that heterophilic Dpr-DIP interactions instruct synaptic specificity through a lock and key-type of mechanism (Carrillo et al., 2015; Tan et al., 2015). Dprs and DIPs have 2 or 3 Ig domains in their extracellular regions, respectively, and Dpr-DIP complexes bear a striking resemblance to the complexes of mammalian IgSF proteins [reviewed in (Zinn and Ozkan, 2017)]. Previous studies in *Drosophila* have shown that Dpr-DIP interactions regulate cell morphogenesis, cell survival or differentiation (Carrillo et al., 2015), and axon-axon fasciculation (Barish et al., 2018). However, whether Dpr-DIP interactions regulate synaptic specificity independent of these developmental processes remains unknown.

To test whether Dpr-DIP interactions play a specific role in establishing synaptic specificity, we have focused on the lamina of the *Drosophila* optic lobe (Fig. 1A-C) which comprises a highly stereotyped cellular and synaptic architecture that has been extensively characterized in electron microscopy (EM) studies (Meinertzhagen and O’Neil, 1991; Rivera-Alba et al., 2011). Within the lamina, the synaptic terminals of photoreceptors R1-R6 (R cells) and the neurites of second-order lamina neurons L1-L5 (L cells) organize into cylindrical modules called cartridges (Fig. 1A and B). Each cartridge receives input from R cells that detect light from the same point in visual space (Braitenburg, 1967), and neighboring cartridges process information from neighboring points in space, so as to establish a retinotopic map. The core of each cartridge primarily comprises the main axon and dendrites of L1 and L2, which are sandwiched in between a ring of R cell axon terminals (Fig. 1B). The main neurites of L3-L5 are located in the cartridge periphery, although L3 sends dendrites into the cartridge core. R cells synapse *en passant* onto L1-L3 dendrites throughout each cartridge, but L1-L3 cells neither synapse reciprocally onto R cells nor synapse with each other (Fig. 1C). Near the base of each cartridge (i.e. the proximal lamina) L4 extends dendrites into the core of its own cartridge (Fig. 1A) and those of two neighbors and forms reciprocal connections with L2 (Fig. 1C). All L cells send axons into the underlying medulla neuropil where they synapse onto specific target cells.

**Figure 1.**
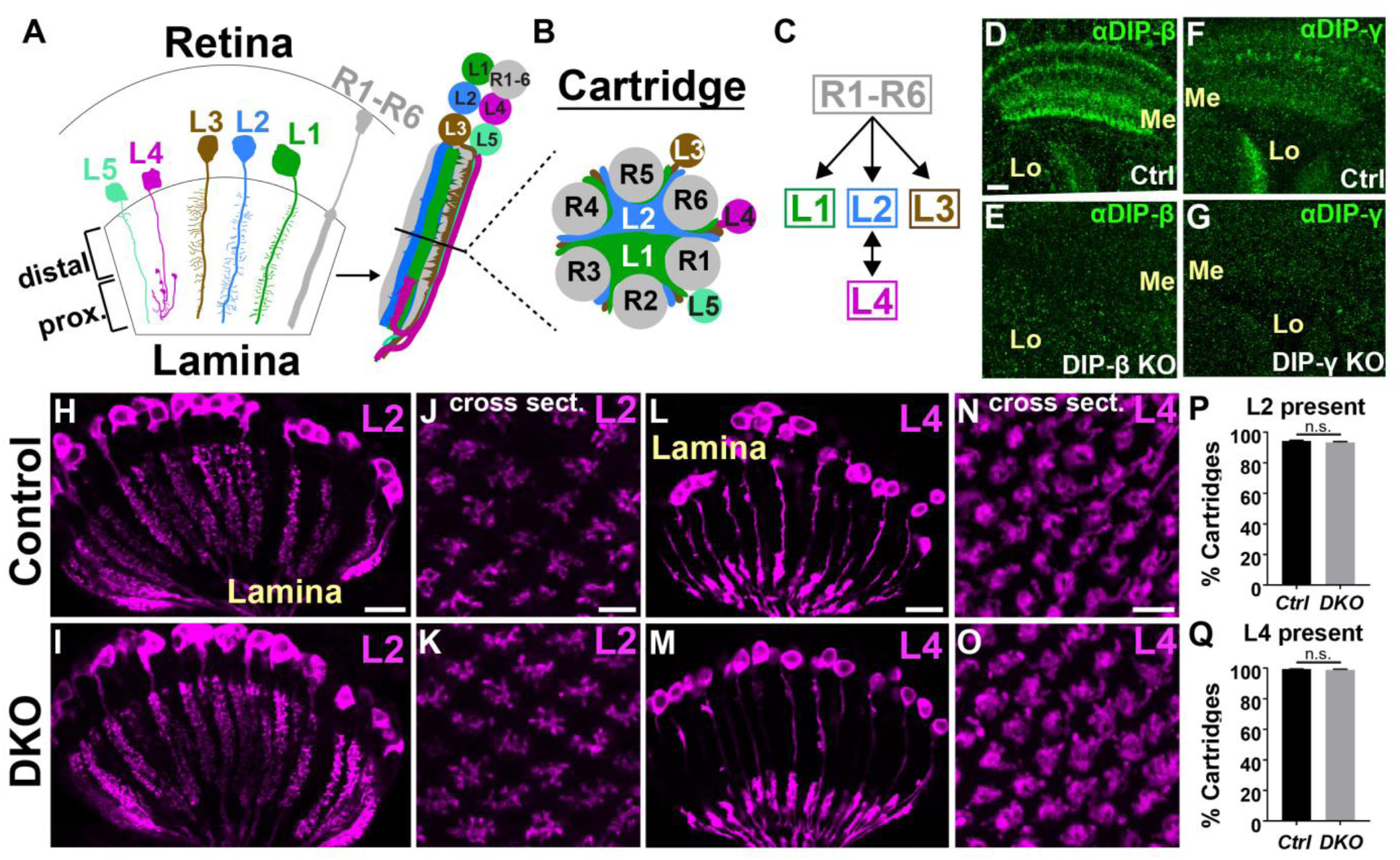
DIP proteins are not required for the normal development of L2 and L4 neurons. (A-C) Cellular and synaptic organization of the lamina. (D-G) Confocal images showing DIP-β (D and E) or DIP-γ (F and G) immunolabeling (green) in the medulla (Me) and lobula (Lo) neuropils at 40-48 hours after puparium formation (h APF). (D and F) In control flies (wild type) DIPs-β and γ are expressed in the medulla and lobula. n=2 brains, scale bar (D)= 10µm. (E and G) The expression of DIPs-β and γ in the medulla and lobula is severely reduced in flies homozygous for *DIP-β* or *γ* null mutations, respectively. n=2 brains per genotype. (H-O) Confocal images of L2 (H-K) or L4 (L-O) neurons in control (H, J, L, N) or DKO (I, K, M, O) flies (1-2 day old adults). (H and I) Longitudinal section of lamina cartridges. L2 neurons (magenta-myristoylated-tandem tomato [myr-tdTOM]) were labeled using a specific driver (see Methods section). L2 neurons in control (n=8 brains) and DKO (n=9 brains) flies were morphologically indistinguishable. Scale bar (H)= 10µm. (J and K) Cross sections through the lamina (single 0.4µm image sections) show that the morphology and spacing of L2 neurites within cartridges are indistinguishable in control (n=9 brains) and DKO flies (n=8 brains). Scale bar (J)= 5µm. (L and M) L4 neurons viewed in their longitudinal axis in the lamina (labeled using a specific driver-see Methods section). The morphologies of L4 neurons in control (n=9 brains) and DKO (n=9 brains) flies were indistinguishable. Scale bar (L)= 10µm. (N and O) Cross sections (13.6 µm maximum projections) through the lamina show that the morphology and spacing of L4 neurons in control (n=9 brains) and DKO (n=7 brains) flies are highly similar. Scale bar (N)= 5µm. (P and Q) Quantification of L2 and L4 cell numbers in control and DKO flies (data are represented as a mean +/- SEM). See methods for description of how cells were counted. Note that not all L2 and L4 neurons are fluorescently labeled in control flies (<100%). (P) The average percentages of lamina cartridges containing fluorescently labeled L2 neurons in control (95%; n=7 brains) and DKO (93%; n=7 brains) flies did not significantly differ. (Q) The average percentages of cartridges containing L4 neurites in control (99%; n=8 brains) and DKO (99%; n=6 brains) flies were identical.

Using loss- and gain-of-function approaches we have exploited the cell-type specificity of synapse formation within lamina cartridges to address whether Dpr-DIP interactions are necessary for synaptic specificity, and sufficient to promote synapse formation between specific cell types. We find that rather than being required for synaptogenesis DIP proteins act to prevent the formation of ectopic synapses, and that mis-expression of DIP proteins is sufficient to promote synapse formation with cell types expressing matching Dprs. These findings suggest that Dpr-DIP interactions regulate synaptic specificity by biasing synapse formation towards specific cell types, thereby preventing promiscuous synapse formation. We consider our findings within the broader context of synapse assembly.

## RESULTS

### DIP proteins are not required for the normal development of L2 and L4 neurons

To test whether Dpr-DIP interactions are necessary for synaptic specificity we focused on L2 and L4 neurons, which selectively form reciprocal connections in the proximal lamina (Meinertzhagen and O’Neil, 1991; Rivera-Alba et al., 2011) (Fig.1C). L2 contacts L1 extensively throughout the cartridge (Fig. 1B), yet is only pre-synaptic in the proximal region where it synapses primarily onto L4, while L4 dendrites extend into the proximal cartridge core (Fig. 1A) and encounter L1 and L2 processes, yet primarily synapse onto L2. Previously, through use of RNA-seq and GAL4 reporters both L2 and L4 were found to express a single DIP during pupal development, L4 expressing DIP-β and L2 expressing DIP-γ (Tan et al., 2015). DIP-β is known to bind to 5 different Dpr proteins in vitro (Dprs 6, 8, 9, 11, 21) (Carrillo et al., 2015; Ozkan et al., 2013), all of which are expressed in L2 neurons (Tan et al., 2015).DIP-γ is known to bind 4 Dprs in vitro (Dprs 11, 15, 16, 17) (Ozkan et al., 2013). While L4 was not found to express any of these Dprs at 40 hours after puparium formation (h APF) (Tan et al., 2015), we reasoned that one or more of these may be expressed in L4 during synapse formation which occurs later in development (see below). Thus, we hypothesized that interactions between DIPs-β, γ and their cognate Dprs may promote selective synapse formation between L2 and L4 neurons. As DIPs-β and γ bind to many Dprs, to test this hypothesis we concentrated our efforts on addressing the functions of DIPs-β and γ.

Using the CRISPR/Cas9 system we generated early stop mutations near the translational start sites of DIPs-β and γ (see Methods section). DIP-β and γ immunolabeling was eliminated in the optic lobes of flies homozygous for these mutations (Fig. 1D-G), demonstrating their efficacy in disrupting DIP function. Before directly testing whether DIPs-β and γ are necessary for synaptic connections between L2 and L4, we first assessed if these proteins are important for the development of these neurons. To accomplish this, we labeled L2 or L4 neurons using cell type-specific GAL4 drivers (Tuthill et al., 2013) in flies doubly heterozygous (control) or doubly homozygous (double knockout [DKO]) for the mutations we generated. The morphologies of L2 and L4 neurons in control and DKO flies were indistinguishable (Fig.1H-O) and disrupting DIP function did not affect cell numbers (Fig. 1P and Q). These findings demonstrate that DIPs-β and γ, and thus Dpr-DIP interactions, are not necessary for the normal development of L2 and L4 neurons.

### DIP function is necessary to prevent the formation of ectopic synapses

Next, we sought to determine whether Dpr-DIP interactions are necessary for selective synapse formation between L2 and L4 neurons. To address this, we used synaptic tagging with recombination (Chen et al., 2014; Peng et al., 2018) to label L2-L4 synapses selectively. Using STaR the active zone protein Bruchpilot (Brp) (Wagh et al., 2006) can be tagged in a cell type-specific manner depending on the expression of Flp recombinase (Golic and Lindquist, 1989), while being expressed from its native promoter within a bacterial artificial chromosome (BAC). Moreover, the cells that express tagged-Brp can also be made to express a fluorescent reporter through the LexA/LexAop system (Lai and Lee, 2006), providing a context in which to assess Brp localization. As L2 and L4 are the only L cells that are pre-synaptic in the lamina and because they predominantly synapse with each other (Meinertzhagen and O’Neil, 1991; Rivera-Alba et al., 2011), selectively expressing Flp in L cells allows selective visualization of L2-L4 synapses in the proximal lamina (Fig. 2A-B’). In the absence of DIP function, we expected to observe a reduction in the number of Brp puncta in the proximal lamina, which would indicate a loss of L2-L4 synapses. However, in DIP DKO flies the number of Brp puncta in the proximal lamina was qualitatively normal compared with control flies (*β*+/+, *γ*+/-) (Fig. 2B-C’), demonstrating that DIP function is not necessary for synapse formation. Interestingly, we observed abnormally high numbers of Brp puncta in the distal lamina of DKO flies compared to controls, indicating that DIP function may be necessary to prevent the formation of ectopic synapses.

**Figure 2.**
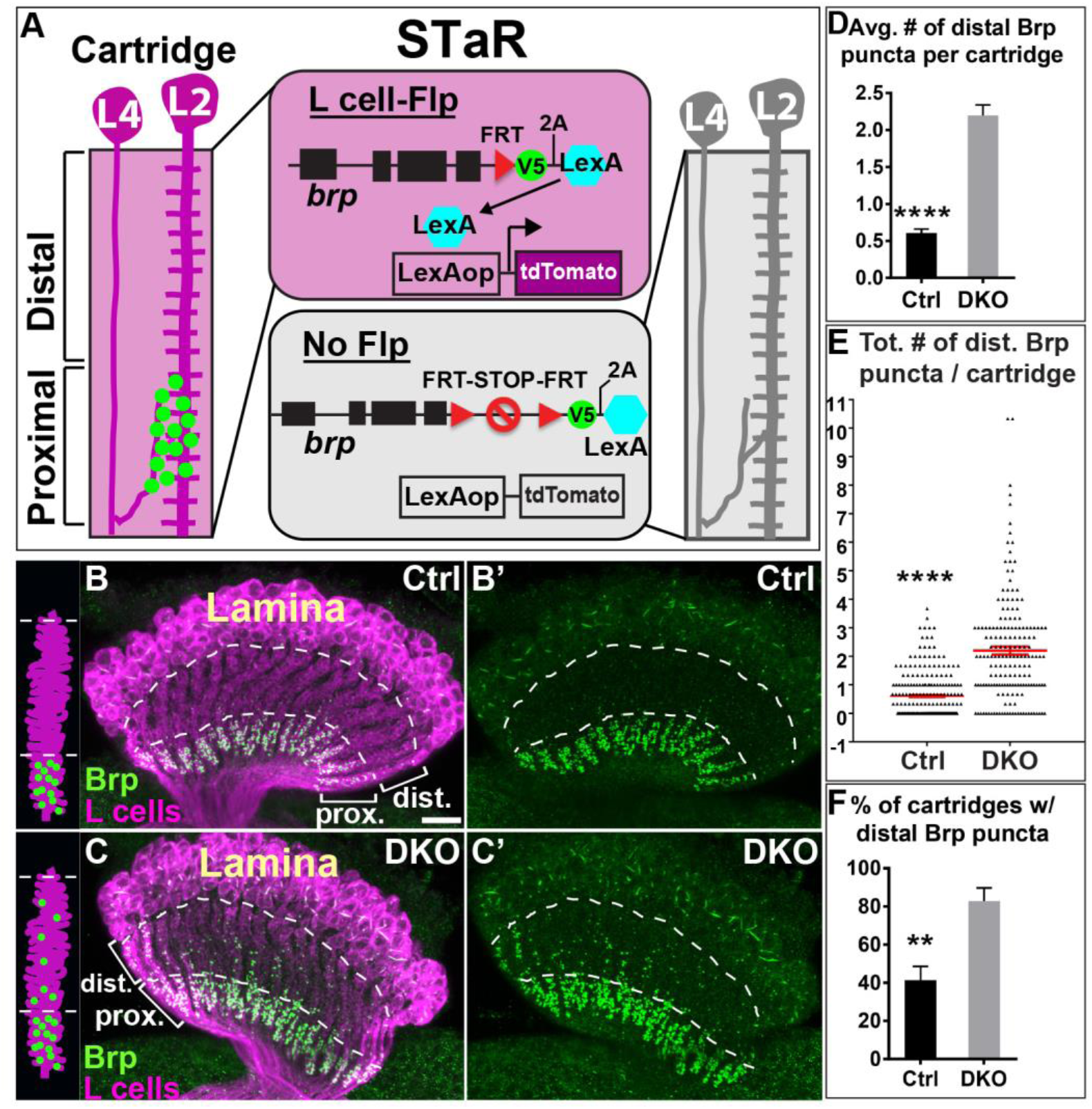
DIP function is necessary to prevent the formation of ectopic synapses. (A) Labeling L2-L4 synapses using STaR. In the absence of Flp recombinase (gray panel) Brp is expressed from its native promoter within a bacterial artificial chromosome, but a transcriptional STOP sequence prevents incorporation of an epitope tag (smFPV5) (Viswanathan et al., 2015) and co-translation of LexA. (Magenta panel) When Flp recombinase is expressed in L cells (27G05-Flp) (Peng et al., 2018) the transcriptional STOP is excised, Brp becomes tagged and LexA is co-translated via the 2A peptide (Ryan and Drew, 1994). LexA then enters the nucleus to drive expression of myr-tdTOM which allows identification of the L cells expressing tagged Brp. As L2 and L4 are the only L cells that are pre-synaptic in the lamina, this allows selective visualization of synapses formed by these neurons. (B-C’) Confocal images (longitudinal plane of the lamina cartridges) showing the distribution of Brp (green-smFPV5) expressed in L cells (magenta-myr-tdTOM) in the laminas of control (*β*+/+, *γ*+/-) or DKO (*β*-/-, *γ*-/-) flies. The area enclosed by the dotted lines indicates the distal lamina (dist.), and the region beneath the most proximal dotted line is the proximal lamina (prox.). (B and B’) In control flies (n= 9 brains), Brp is restricted to the proximal lamina where L2 and L4 neurons are known to form synapses. Scale bar (B)= 10µm. (C and C’) In DKO flies (n= 7 brains), Brp is still localized to the proximal lamina, but ectopic Brp puncta are present in the distal lamina. (D) The average number of Brp puncta in the distal halves of lamina cartridges in control (n= 225 cartridges; 9 brains) and DKO (n= 175 cartridges; 7 brains) flies. Data are represented as a mean +/- SEM. (See also Figure S1) (E) Total number of Brp puncta in the distal halves of cartridges in control (n= 225 cartridges) and DKO (n=175 cartridges) flies. Each triangle represents a cartridge. Red bar indicates +/- SEM. (F) Shows the percentages of cartridges containing distal Brp puncta in control (n= 9 brains) and DKO (n= 7 brains) flies. Data are represented as a mean +/- SEM.

To quantify this phenotype, we imaged down the long axis of lamina cartridges using confocal microscopy and took Z-stacks of the laminas of control and DKO flies. This allowed us to clearly visualize individual cartridges in the lamina clearly and count the number of Brp puncta in their distal halves. In total, we scored 225 control cartridges (9 brains) and 175 DKO cartridges (7 brains). We found that the average number of Brp puncta in the distal region of cartridges was significantly higher in DKO versus control flies (Fig. 2D), as was the percentage of cartridges containing distal Brp puncta (Fig. 2F). In addition, when cartridges did contain distal Brp puncta, cartridges from DKO flies contained significantly more puncta (Fig. 2E). Taken together, our loss-of-function studies demonstrate that DIP function is necessary for proper synaptic connectivity independent of cell survival and morphology.Moreover, our results suggest that rather than being required for synapse formation, DIP proteins are necessary to prevent the formation of ectopic synapses. As L4 processes are restricted to the proximal lamina, ectopic synapses in the distal lamina likely represent L2 synapses onto other cells in the cartridge.

### In the absence of DIP function ectopic synapses form through abnormal synapse formation

We reasoned that ectopic synapses observed in DKO flies could result from abnormal patterns of synapse formation or a deficit in synapse refinement. To address this, we used STaR to label Brp in L cells (Fig. 2A) and assessed Brp localization during development in the lamina of wild type flies. We hypothesized that if ectopic synapses are caused by defects in refinement, then at some point during development we should observe Brp puncta in the distal lamina that are then lost at later stages. Conversely, if ectopic synapses in DKO flies result from abnormal synapse formation, then at no point during development should we observe significant numbers of Brp puncta in the distal lamina. Our experiments revealed the latter to be the case. We found that L2-L4 synapses form at 70h APF, and at no time thereafter did we observe significant numbers of Brp puncta in the distal lamina (Fig. 3A-D’). Thus, ectopic synapses formed in the absence of DIP function result from abnormal synapse formation.

**Figure 3.**
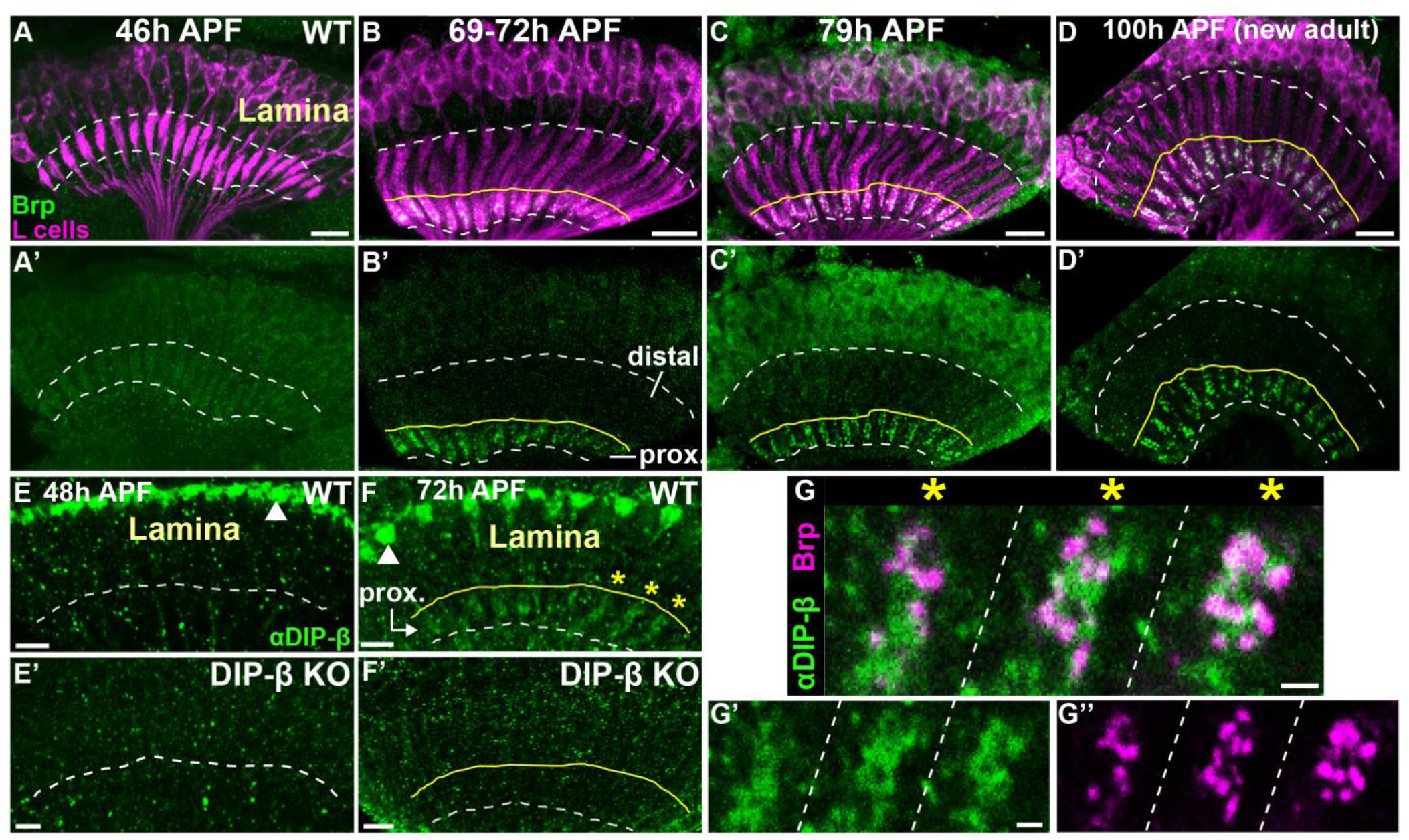
Disrupting DIP function causes abnormal synapse formation and DIP-β localizes to the proximal lamina during the formation of L2-L4 synapses. (A-D’) Confocal images in a longitudinal plane of lamina cartridges showing the developmental timing of Brp expression (green-smFPV5) in L cells (magenta-myr-tdTOM) in the lamina. h APF= hours after puparium formation. The dotted lines delineate the lamina neuropil. The area between the distal dotted line and the solid yellow line delineates the distal lamina. The region between the solid yellow line and the proximal dotted line delineates the proximal lamina. The proximal lamina is defined here by the appearance of synapses in L2 and L4 neurons (69-72h APF). n=2 brains (46h APF; 79h APF), n=3 brains (69-72h APF; 100h APF). Scale bars (A-D)= 10µm. (E-F’) DIP-β immunolabeling (green) in the laminas of wild type and *DIP-β* KO (*β*-/-) flies during pupal development. The dotted white line indicates the proximal edge of the lamina neuropil. The area between the solid yellow line and the dotted white line (F and F’) shows the proximal lamina (prox.) (as in B and B’). White arrowheads (E and F) show DIP-β expression in non-L4 neurons likely to be LaWF2 neurons (Tuthill et al., 2013). Scale bars= 5µm. (E and E’) At 48h APF DIP-β expression in the lamina is only detected in the very distal region of the neuropil, most likely corresponding to LaWF2 neurons. (WT, n= 2 brains) (*DIP-β* KO, n= 2 brains). (F and F’) At 72h APF DIP-β immunolabeling is observed in the proximal lamina, and this labeling is eliminated in *DIP-β* KO flies. Yellow stars show DIP-β expression in the proximal regions of 3 cartridges shown at higher magnification in G-G’’. (WT, n= 6 brains) (*DIP-β* KO, n= 2 brains). (G-G’’) Co-labeling of DIP-β (green) and Brp (magenta-smFPV5) in the proximal regions of 3 lamina cartridges (yellow stars in F) separated by dotted lines. Scale bars (G and G’)= 1µm.

### DIP-β localizes to the proximal lamina during the formation of L2-L4 synapses

To gain insight into how DIP proteins antagonize ectopic synapse formation we assessed the subcellular localization of DIPs-β and γ in the lamina during pupal development through immunolabeling and confocal microscopy. Strong DIP-β immunolabeling was detected in the most distal region of the lamina neuropil throughout pupal development (Fig. 3E and F, white arrowhead). This labeling is not consistent with expression in L4 and likely represents DIP-β expressed in LaWF2 neurons (Tuthill et al., 2013). Outside this labeling pattern, we did not detect DIP-β protein in the lamina at 24 (not shown) or 48h APF (Fig. 3E and E’) but we did observe labeling in the proximal lamina at 72h APF (Fig. 3F), around the time of L2-L4 synapse formation (Fig. 3B and B’). This labeling is consistent with DIP-β localization to L4 dendrites and was eliminated in flies homozygous for the *DIP-β* null mutation (Fig. 3F’). To shed light on whether DIP-β localizes to developing synapses, we simultaneously visualized DIP-β in the proximal lamina (immunolabeling) and Brp in L2 and L4 neurons (STaR, Fig. 2A) at 72h APF using confocal microscopy (Fig. 3G-G’’). We found that DIP-β was more diffusely distributed within the cartridge than Brp, but that some of the DIP-β protein appeared to be organized into clusters that overlapped with or were adjacent to Brp puncta. This was consistent between cartridges and across brains. Taken together, these findings show that DIP-β localizes to the proximal lamina during the onset of synapse formation in L2 and L4 neurons. Moreover, our data are consistent with a subset of DIP-β protein localizing to developing synapses. We were unable to detect DIP-γ protein in the lamina at any stage of pupal development (not shown). However, it’s possible that DIP-γ is expressed at low levels in L2 neurons and that DIP-γ immunolabeling is not sufficiently sensitive to detect this expression.

Collectively, our loss of function, developmental and protein localization studies show that DIP-β localizes to the proximal lamina during L2-L4 synapse formation but that DIP function is not necessary for synaptogenesis, and rather acts to prevent ectopic synapse formation (see Discussion).

### DIP mis-expression is sufficient to change the synaptic connectivity of L cells in a predictable manner

While our experiments exploring DIP function in L2 and L4 neurons indicate that Dpr-DIP interactions are not required for synapse formation in the examined context, we reasoned that Dpr-DIP interactions may be sufficient to promote synapses to form between specific cell types. To test this possibility, we exploited the cell type-specificity of synaptic connections between L cells and R cells in the lamina. Within each cartridge R cells synapse *en passant* onto L1-L3, but L1-L3 do not reciprocally synapse onto R cells, nor do they synapse with each other (Meinertzhagen and O’Neil, 1991; Rivera-Alba et al., 2011) (Fig.1C). This specificity is quite striking given that processes of these neurons are densely packed within the cartridge and contact each other extensively. L cells express high levels of Dprs (Tan et al., 2015) and in general, both L cells and R cells express low levels of DIPs (Tan et al., 2015; Zhang et al., 2016). Thus, we hypothesized that if Dpr-DIP interactions promote synapse formation then mis-expressing DIPs in R cells should cause L cells to synapse onto R cells. Likewise, mis-expressing DIPs in L1-L3 should cause these cells to synapse with each other.

To test this hypothesis, we mis-expressed DIPs-γ and ε either together or independently in R cells and visualized L cell synapses in the lamina using STaR as in the DIP KO experiments. We chose these DIPs because they have broad Dpr binding specificities and are known to bind to Dprs expressed in L cells (Carrillo et al., 2015; Ozkan et al., 2013; Tan et al., 2015). In control flies, L cell synapses were restricted to the proximal lamina where L2 and L4 form reciprocal connections (Fig. 4A and A’).Strikingly, mis-expression of both DIPs-γ and ε (Fig. S1A and A’) or DIP-γ alone (Fig. 4B and B’) caused L cells to form streams of ectopic synapses throughout lamina cartridges. On average, 34% of lamina cartridges in adult (1-2 day old) flies mis-expressing DIP-γ in R cells displayed clusters of ectopic L cell synapses in the distal lamina, while none of the cartridges in control flies showed this phenotype (Fig.4D). Mis-expression of DIP-ε alone did not cause the ectopic synapses to form (Figs. S1B and B’). As both DIPs-γ and ε were strongly expressed in R cells in mis-expression experiments (Fig. S1C and D) the differences in their abilities to promote ectopic synapse formation may reflect alternative functions for these proteins (see Discussion). Cross sections through the lamina of flies mis-expressing DIP-γ in R cells showed that, in general, the organization of L cell processes within cartridges and cartridge spacing both appeared normal (Fig. S1E and F). In addition, this revealed that ectopic synapses frequently formed on the edges of L cell profiles consistent with the positions of R cell axon terminals (Fig. 4C). Mis-expression of DIPs-γ and ε in L cells also caused L cells to form ectopic synapses throughout lamina cartridges (Fig. S1G-H’). Together, these findings show that DIP-mis-expression causes L cells to form ectopic synapses in a predictable manner.

**Figure 4.**
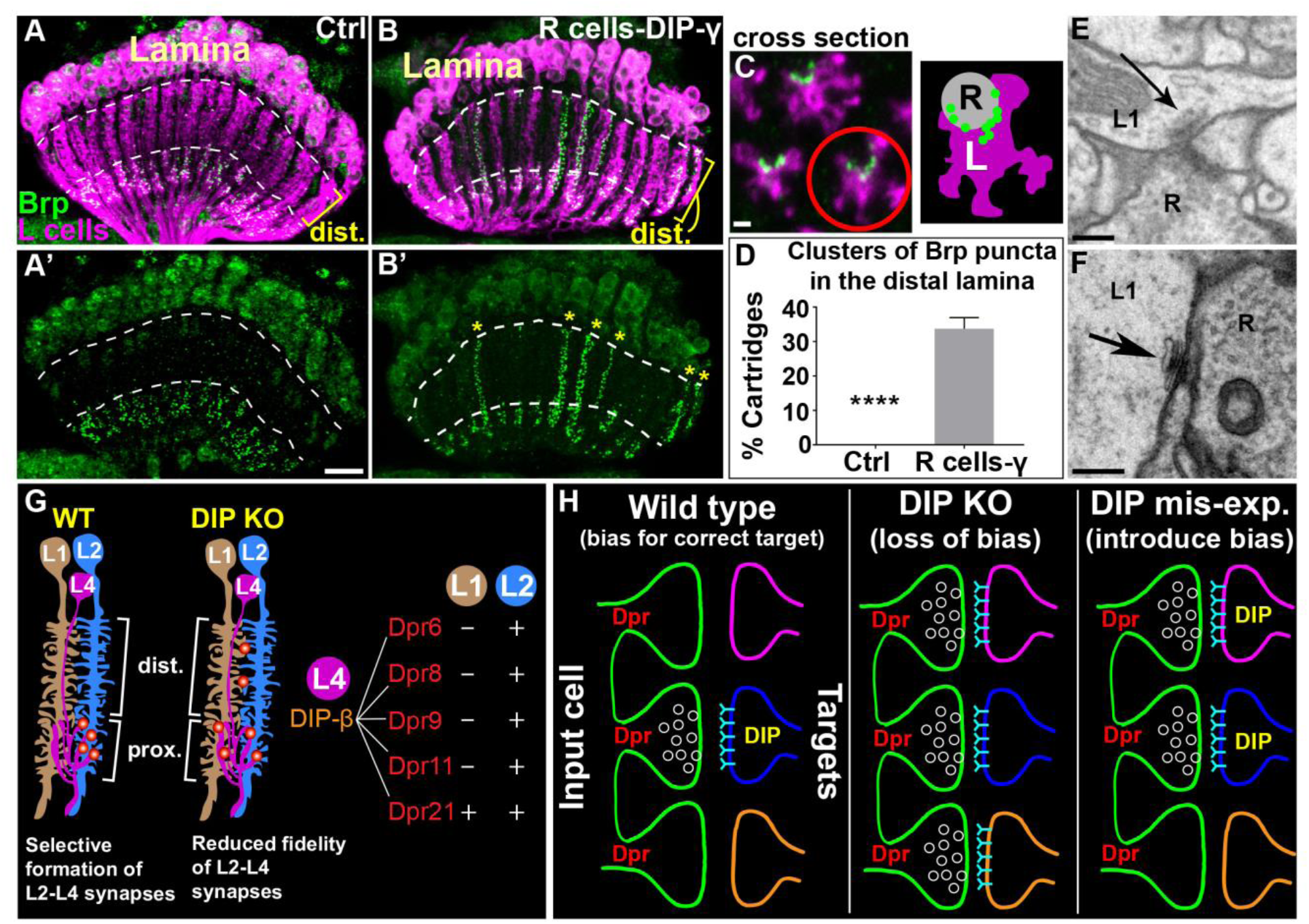
DIP mis-expression is sufficient to change the synaptic connectivity of L cells in a predictable manner. (See also Figures S1 and S2, and Table S1) (A-B’) Confocal images (longitudinal plane of lamina cartridges, 1-2 day old adults) showing the distribution of Brp (green-smFPV5) expressed in L cells (magenta-myr-tdTOM) in the laminas of control (UAS-DIP-γ or GMR-GAL4 alone) flies or flies expressing DIP-γ in R cells (UAS-DIP-γ and GMR-GAL4). The area between the dotted white lines delineates the distal lamina (dist.). Scale bar (A’)= 10µm. (A and A’) Brp is restricted to the proximal lamina in control flies (region beneath the most proximal white dotted line). (B and B’) In flies mis-expressing DIP-γ in R cells streams of Brp puncta are detected throughout lamina cartridges (yellow stars in B’). (c) (Left panel) A confocal image of a cross section through the lamina of a fly mis-expressing DIP-γ in R cells. (Right panel) A schematic of the cartridge in the left panel that is circled in red. L indicates L cell neurites within the cartridge (magenta), and R indicates a potential R cell terminal that is post-synaptic to L cells in the cartridge (green dots= L cell synapses). (D) Quantification of the percentages of cartridges containing clusters of Brp puncta in the distal lamina in control flies (n= 11) and flies mis-expressing DIP-γ in R cells (n= 11). Data are represented as a mean +/- SEM. (E and F) Putative L1-R cell synapses in flies mis-expressing DIPs-γ and ε in R cells identified by EM. Scale bars= 200nm. (G) Working model of how Dpr-DIP interactions regulate selective synapse formation between L2 and L4 neurons. In wild type flies, L2 and L4 selectively synapse with each other in the proximal lamina. In DIP KO flies, we propose that the fidelity of L2-L4 synapse formation is reduced resulting in synapse formation with additional cell types. In the distal lamina, we speculate that this manifests as ectopic synapses made by L2 onto cell types other than L4 (e.g. L1). And in the proximal lamina, we hypothesize that L2 and L4 still form synapses, but also synapse with inappropriate partners (e.g. L1). The preference of L2 and L4 to synapse with each other over L1 neurons may be due to the differential expression of Dprs that bind DIP-β in L2 and L1 neurons [reflects Dpr expression in L1 and L2 at 40h APF (Tan et al., 2015)]. (H) Dpr-DIP interactions may regulate synaptic specificity by biasing synapse formation towards specific cell types. (Left panel) In a wild type background, Dpr-DIP interactions bias synapses to form between specific cell types, potentially by concentrating synaptic machinery to specific cell-cell contacts. (Middle panel) When Dpr-DIP interactions are disrupted there is a reduced bias for the correct synaptic partner, resulting in promiscuous synapse formation with inappropriate partners. (Right panel) Inducing ectopic Dpr-DIP interactions introduces a bias for inappropriate synaptic partners.

To visualize the morphologies and cellular composition of ectopic synapses induced by DIP mis-expression we utilized electron microscopy. We cut the lamina of a fly mis-expressing DIPs-γ and ε in R cells into 50-60 nm sections and imaged these using transmission electron microscopy (TEM). We then identified the synapses formed in the sections and assigned them to cellular profiles based on previously established criteria (Meinertzhagen, 1996; Meinertzhagen and O’Neil, 1991). We found that the positions of cell types within the cartridge were normal, with L1 and L2 always together in the axis surrounded by R cell terminals (Fig. S2). In addition, the numbers of synapses formed by R cells (R cells pre-synaptic) was similar to those reported previously, with the 6 R cell profiles together contributing 330 synapses (Table S1) (Meinertzhagen and O’Neil, 1991; Rivera-Alba et al., 2011).Thus, DIP-mis-expression did not significantly perturb the general cellular architecture of the cartridge, or synapse formation in R cells. We identified 86 L cell synapses within the cartridge (L cells pre-synaptic) (Table S1), ∼3-4 times more L cell synapses than was previously reported for wild type cartridges (Meinertzhagen and O’Neil, 1991; Rivera-Alba et al., 2011). These were distributed throughout the cartridge, with 31 L cell synapses in the distal half (Table S1). In addition, we identified synapses formed by L1 (x12), L3 (x13), and L5 (x5) neurons which were previously not found to be pre-synaptic in the lamina (we also identified L2 and L4 synapses). In some cases, identified R cell profiles were adjacent to L cell presynaptic sites (Fig. 4E and F) consistent with L cell to R cell synapses, although a full reconstruction would be necessary to determine the degree to which L cell synapses form onto R cells upon DIP mis-expression. Together, these findings complement and support our confocal analyses and show that DIP-mis-expression promotes synapse formation in a manner predicted by Dpr expression.

Collectively, our confocal and EM phenotypic analyses of DIP mis-expression indicate that Dpr-DIP interactions are sufficient to promote synapse formation between specific cell types, and are consistent with Dpr-DIP interactions acting instructively to control synaptic specificity.

## DISCUSSION

Neurobiologists have long thought that appropriate synaptic partners express complementary molecules that allow them to identify each other within a dense meshwork of alternative neurites through a lock-and-key mechanism. A common interpretation of this idea is that interactions between complementary adhesion or recognition molecules are necessary for synaptogenesis. Based on their heterophilic binding and matching expression in synaptically connected cell types, Dpr and DIP IgSF proteins have been proposed to play an instructive role in regulating synaptic specificity through a complementary binding mechanism. The findings we present here support this hypothesis. However, rather than being necessary for synaptogenesis, we propose that Dpr-DIP interactions regulate synaptic specificity by biasing synapses to form between specific cell types (see below). In this view, synaptic specificity is a bias rather than an absolute or categorical recognition, and the precision of synaptic connectivity is established by limiting promiscuous synapse formation.

### Dpr-DIP interactions may regulate synaptic specificity by biasing synapse formation towards specific cell types

Overall, our findings support the idea that Dpr-DIP interactions play an instructive role in establishing synaptic specificity. We hypothesize that interactions between L2 and L4 neurons mediated by DIP-β, γ, or both, and their cognate Dprs promote the formation of L2-L4 synapses by concentrating synaptic machinery at sites of mutual contact. Disrupting these interactions then causes, we propose, synaptic machinery to be diffusely distributed within these cells resulting in either randomized synapse formation or a broader choice of synaptic targets (Fig. 4G). We speculate that in the distal lamina of DIP KO flies, L2 forms ectopic synapses with cell types other than L4, and in the proximal lamina L2 and L4 still synapse with each other but also form connections less discriminately with incorrect partners. Thus, our model proposes that when Dpr-DIP interactions are disrupted synapses still form but reflect a loss of bias for the correct partners, and that inducing ectopic Dpr-DIP interactions (e.g. through DIP mis-expression) introduces bias and promotes synapses to form between incorrect partners (e.g. L-R or L-L synapses) (Fig. 4H). This is reminiscent of how the proteins SYG-1 and SYG-2 are thought to regulate synaptic specificity in *C. elegans*. SYG-1 expressed in the HSNL neuron binds to SYG-2 expressed in vulval epithelial cells (guidepost cells), to restrict the subcellular location of synapse formation, biasing HSNL to synapse with specific partners (Shen and Bargmann, 2003; Shen et al., 2004). An important difference between the SYG proteins and Dprs/DIPs is that Dpr-DIP interactions occur between synaptic partners rather than with guidepost cells. As Dpr-DIP complexes are highly similar to complexes of mammalian IgSF proteins (Zinn and Ozkan, 2017), we speculate that the model we propose may represent a widespread strategy for establishing synaptic specificity.

### DIP function is necessary to prevent ectopic synapse formation

The localization of DIP-β to the proximal lamina during the stage of L2-L4 synapse formation and the abnormal formation of synapses in the distal lamina in the absence of DIP function both suggest that DIP-β, and potentially DIP-γ, play a role in restricting synaptic machinery to sites of L2-L4 contacts. Analysis of synapse formation in flies in which DIP-β or γ are selectively disrupted in L4 or L2, respectively, will address the contributions of these proteins to synaptic connectivity and their cellular requirements.

DIPs-β and γ are likely to function through binding in *trans* with cognate Dprs expressed respectively in L2 or L4 neurons (Fig. 4G). At 40h APF, L2 neurons express all 5 Dprs known to bind to DIP-β (Dprs 6, 8, 9, 11, 21) and also express both of the Dprs within this group (Dprs 6, 11) that were tested for cellular expression at 72h APF (Tan et al., 2015), when L2-L4 synapses begin to form (Fig. 3B and B’). By contrast, L1 neurons, which along with L2 represent the bulk of the cartridge core, express just one of these Dprs (Dpr21) at 40h APF and neither of the Dprs tested at 72h APF (Tan et al., 2015). Based on this differential Dpr expression it is likely that DIP-β preferentially binds with L2-Dprs over those expressed in L1, and we speculate that this is important for selective L2-L4 synapse formation in the lamina. With respect to DIP-γ, L1-L3 express cognate Dprs (Dprs 11, 15, 16, 17) at 40h APF, and L2 also expresses Dpr11 at 72h APF. However, Dprs 15 and 17 are not expressed by L cells at 72h APF (Tan et al., 2015). The cellular expression of Dpr16 at 72h APF has not been addressed, and it may be expressed in L4 at that time. Thus, it is possible that interactions between DIP-γ expressed in L2 and Dpr16 in L4 are also important for the selective formation of synapses between L2 and L4 neurons. Determining how disrupting Dpr function in L2 and L4 neurons or changing the Dpr binding specificities of DIPs-β and γ affects synapse formation will test these hypotheses. Additionally, it will be important to determine if Dpr and DIP proteins cluster together at developing L2-L4 synapses and interact with synaptic proteins. Given that Dprs and DIPs lack obvious intracellular signaling motifs and that many are predicted to be linked to the plasma membrane through a lipid anchor, if they interact with synaptic machinery this would be likely to occur through co-receptors. Thus, identifying such co-receptors may be the key to understanding how Dpr-DIP interactions regulate synaptic specificity.

As L2 and L4 are the only L cells that are pre-synaptic in the lamina, they are the L cells most likely to form ectopic synapses in the distal lamina in DIP KO flies. L4 processes are largely restricted to the proximal region of each cartridge, and we hypothesize that ectopic synapses therefore represent L2 synapses onto other cell types in the cartridge (e.g. L1) (Fig. 4G). With respect to the proximal lamina, if target selection becomes randomized, or broader, in the absence of DIP function as we propose in our model then we expect that L2 and L4 still form synapses in this region but also synapse with inappropriate partners. To test this idea, it is necessary to identify the cellular components of L cell synapses in the distal and proximal lamina in DIP KO flies. This could potentially be accomplished by co-labeling pre- and postsynaptic proteins expressed in different cell types in the lamina using STaR and cell-specific drivers, or through EM studies.

### DIP mis-expression is sufficient to alter synaptic connectivity in a predictable manner

If Dpr-DIP interactions act instructively to control synaptic specificity, as is suggested by their complementary expression in synaptic partners, then they should be sufficient to promote synapse formation between specific cell types. Our mis-expression experiments support that this is indeed the case. We hypothesize that mis-expressing DIP-γ in R cells or L cells promotes *trans* interactions between DIP-γ and cognate Dprs expressed in L cells that recruit synaptic machinery resulting in synapse formation. Eliminating the ability of DIP-γ to bind Dprs or disrupting the function of Dprs that bind DIP-γ in L cells will determine whether specific Dpr-DIP interactions contribute to ectopic synapse formation in mis-expression experiments.

It is interesting that ectopic L cell synapses were not induced by mis-expressing DIP-ε, which is known to bind to 7 Dprs (Ozkan et al., 2013) (more than any other DIP), all of which are expressed in L cells during pupal development based on RNA-seq analyses (3 of these were confirmed at the protein level) (Tan et al., 2015). One possibility is that different Dpr-DIP interactions support different functions. Indeed, DIP-γ has alternative functions at the NMJ and in the medulla (Carrillo et al., 2015) indicating that even the same DIPs can function differently in different contexts. Alternatively, it is possible that Dprs that bind DIP-ε are not expressed at sufficient levels in L cells, or that most of these proteins localize to L cell axons in the medulla and are not available for binding. One way of distinguishing between these possibilities is to mis-express hybrid versions of DIPs-ε and γ and assess their ability to induce ectopic synapse formation. These could include versions with switched Dpr binding specificities and versions in which Dpr binding remains intact but other regions are switched.

In this study, we have manipulated DIP expression in multiple cell types simultaneously as a proof of principle to show that such manipulations can predictably change synaptic connectivity. An exciting next step is to determine if more precise (i.e. cell type-specific) manipulations of Dpr-DIP interactions can alter connectivity and behavior in a predictable manner.

### Model of synapse assembly

Synapse formation is necessarily robust. In the absence of the correct synaptic targets neurons have the capacity to synapse with other cell types. Along the same lines, our findings indicate that molecules that mediate synaptic specificity are not necessary for synaptogenesis. This argues against the idea that interactions between appropriate synaptic partners initiate synapse assembly and rather suggests that synaptogenesis occurs independently in a largely indiscriminate manner, with specificity molecules introducing bias to the process. In organisms with highly primitive nervous systems the developmental order of neuron birth and targeting events such as axon guidance may have been sufficient to establish the precision of synaptic connectivity without including restraints on synaptogenesis. And in more advanced organisms with complex nervous systems cell recognition mechanisms may have been adapted to overlay specificity onto promiscuous synapse formation. Moving forward, a key question to be addressed is how molecules that mediate specificity communicate with synaptic machinery to establish synapses of the right type with the correct partners.

## ACKNOWLEDGEMENTS

We would like to thank Dr. Larry Zipursky for generously sharing reagents, and Drs. Ginty and Kaeser for many helpful discussions. This research was supported by the NIH/NINDS (National Institute of Neurological Disorders and Stroke) grant K01 K01NS094545 (M.Y.P.), and by the McKnight Foundation (Scholar Award) (M.Y.P.). C.X. was supported by an Alice and Joseph Brooks Postdoctoral Fellowship, and an Edward R. and Anne G. Lefler Center Postdoctoral Fellowship.

## AUTHOR CONTRIBUTIONS

Conceptualization, M.Y.P. and C.X.; Methodology, M.Y.P. and C.X.; Validation, C.X., E.T., E.R., and B.S.; Formal Analysis, M.Y.P., C.X., E.T., D.T., J.B., and I.A.M.; Investigation, C.X., E.T., E.R., and D.T; Resources, M.Y.P., I.A.M., J.P., L.T., and M.C.; Writing-Original Draft, M.Y.P.; Writing-Review and Editing, M.Y.P., C.X., E.T., and I.A.M.; Visualization, M.Y.P., C.X., E.T.; Supervision, M.Y.P., C.X.; Project Administration, M.Y.P.; Funding Acquisition, M.Y.P.

## MATERIALS AND METHODS

### Fly strains

Flies were raised on standard cornmeal-agar based medium. Male and females flies were used at the following development stages: 24, 48, and 72 hr after pupariam formation (h APF), and newly eclosed adults (within 5 hr of eclosion).

The following fly stocks were used:

*53G02AD (II) 29G11DBD (III)* [L2-split-GAL4 (Tuthill et al., 2013)] (gift from the Janelia Research Campus), *79C23S-GS-FRT-stop-FRT-smFPV5-2A-LexAVP16* (Peng et al., 2018), *LexAop-myr::tdTomato (III)* (gift from S.L. Zipursky), *UAS-Flp (II)* (BDSC 4540), *31C06AD (II) 34G07DBD (III)* [L4 split-GAL4 (Tuthill et al., 2013)] (gift from the Janelia Research Campus), *27G05-FLPG5.PEST (attp5)* (gift from the Janelia research campus) (Peng et al., 2018), *GMR-GAL4 (III)* (BDSD 8121), *27G05-GAL4 (attp2)* (BDSC 48073)

### Fly Genotypes

*Figure 1:*

D,F: w; Bl/Cyo; TM2/TM6B

E: w; Bl/Cyo; *DIP-γ*^*1-67*^

G: DIP-β^1-95^; +/+; +/+

H, J: *DIP-β*^*1-95*^ (w, or Y)/w; 53G02AD/UAS-FLP, 79C23S-GS-FRT-stop-FRT-smFPV5-2A-LexAVP16;29G11DBD/ *DIP-γ*^*1-67*^, LexAop-myr::tdTomato

L, N: *DIP-β*^*1-95*^/w; +/UAS-FLP, 79C23S-GS-FRT-stop-FRT-GFPv5-2A-LexAVP16; 31C06AD,34G07DBD/ *DIP-γ*^*1-67*^, LexAop-myr::tdTomato

I, K: *DIP-β*^*1-95*^; 53G02AD/UAS-FLP, 79C23S-GS-FRT-stop-FRT-GFPv5-2A-LexAVP16; 29G11DBD,*DIP-γ*^*1-67*^/ *DIP-γ*^*1-67*^, LexAop-myr::tdTomato

M, O: *DIP-β*^*1-95*^; +/UAS-FLP, 79C23S-GS-FRT-stop-FRT-GFPv5-2A-LexAVP16; 31C06AD, 34G07DBD, *DIP-γ*^*1-67*^/ *DIP-γ*^*1-67*^, LexAop-myr::tdTomato

*Figure 2:*

B, B’: w; 79C23S-GS-FRT-stop-FRT-GFPv5-2A-LexAVP16/ 27G05FLP; *DIP-γ*^*1-67*^, LexAop-myr-tdTOM/TM2(TM6B)

C, C’: *DIP-β*^*1-95*^, 79C23S-GS-FRT-stop-FRT-GFPv5-2A-LexAVP16/ 27G05FLP; *DIP-γ*^*1-67*^, LexAop-myr-tdTOM/ *DIP-γ*^*1-67*^

*Figure 3:*

A-D’: w; Bl(Cyo)/27G05FLP; 79C23S-GS-FRT-stop-FRT-GFPv5-2A-LexAVP16, LexAop-myr-tdTOM/ TM2

E: w; Bl/Cyo; TM2/TM6B

E’, F’: *DIP-β*^*1-95*^; +/+; +/+

F, G, G’, G’’: w; 27G05FLP, 79C23S-GS-FRT-stop-FRT-GFPv5-2A-LexAVP16; TM2/TM6B

*Figure 4:*

A, A’: w; UAS-DIP-γ/27G05FLP; 79C23S-GS-FRT-stop-FRT-GFPv5-2A-LexAVP16, LexAop-myr-tdTOM/TM2(TM6B)

B, B’, C: w; UAS-DIP-γ/27G05FLP; 79C23S-GS-FRT-stop-FRT-GFPv5-2A-LexAVP16, LexAop-myr-tdTOM/GMR-GAL4

E, F: w; UAS-DIP-γ, UAS-DIP-ε/27G05FLP; 79C23S-GS-FRT-stop-FRT-GFPv5-2A-LexAVP16,

LexAop-myr-tdTOM/GMR-GAL4

*Figure S1:*

A, A’: w; UAS-DIP-γ, UAS-DIP-ε/27G05FLP; 79C23S-GS-FRT-stop-FRT-GFPv5-2A-LexAVP16,LexAop-myr-tdTOM/GMR-GAL4

B, B’: w; UAS-DIP-ε/27G05FLP; 79C23S-GS-FRT-stop-FRT-GFPv5-2A-LexAVP16, LexAop-myr-tdTOM/GMR-GAL4

C, D: w; UAS-DIP-γ, UAS-DIP-ε/27G05FLP; 79C23S-GS-FRT-stop-FRT-GFPv5-2A-LexAVP16,LexAop-myr-tdTOM/GMR-GAL4

E: w; Bl (Cyo)/27G05FLP; GMR-GAL4/79C23S-GS-FRT-stop-FRT-GFPv5-2A-LexAVP16, LexAop-myr-tdTOM/TM2(TM6B)

F: w; UAS-DIP-γ/27G05FLP; 79C23S-GS-FRT-stop-FRT-GFPv5-2A-LexAVP16, LexAop-myr-tdTOM/GMR-GAL4

G, G’: w; Bl(Cyo)/27G05FLP; 79C23S-GS-FRT-stop-FRT-GFPv5-2A-LexAVP16, LexAop-myr-tdTOM/27G05-GAL4

H, H’: w; UAS-DIP-r/27G05FLP; 79C23S-GS-FRT-stop-FRT-GFPv5-2A-LexAVP16, LexAop-myr-tdTOM/27G05-GAL4

*Figure S2:*

w; UAS-DIP-γ, UAS-DIP-ε/27G05FLP; 79C23S-GS-FRT-stop-FRT-GFPv5-2A-LexAVP16, LexAop-myr-tdTOM/GMR-GAL4

### Generation of DIP mutants

The method to generate null alleles for *DIP-β, DIP-γ* and *dpr11* via the CRISPR/Cas9 system is described in Xu et al., (submitted to Neuron). Briefly, we chose two protospacer sequences that are ∼50-200 bp away in the genome, in the upstream exons, to create a short deletion causing a nonsense mutation close to the upstream of the protein sequence. High score protospacer sequence was chosen on http://crispr.dfci.harvard.edu/SSC/. We cloned each protospacer into pU6-2-sgRNA-short (Addgene 41700) plasmid and co-injected two plasmids into vas-Cas9 line (BDSC 51323 or 51324, depending on which chromosome the gene is at) in Bestgene. Injected larvae were crossed with double-balanced lines, and screened in F1for single flies carrying the mutation using PCR of the genomic region overlapping with the region to delete. A mutant stock was established from this single F1. Detailed protocols are available upon request.

*Information for CRISPR null mutants:*

*DIP-β*^*1-95*^ deleted sequence: AGCCGGACTTTGTGATTCCGCTGGAGAACGTGACCATCGCCCAAGG

DIP-γ^*1-67*^ deleted sequence:

GGCAGGACGCGAGGCCATCCTGGCCTGCTCGGTGCGCAATCTCGGCAAGAATAAGGTGAGCTAGAATGATTTACCTTGCATTGCAATATATATAATATGATATATAATCCCCTGATAATAGGTTGGTTGGCTGAGAGCCTCCGATCAGACCGTTTTAGCTCTCCAAGGTCGCGTTGTCACCCATAATGCGAGA

Insertion right upstream of the deleted sequence:

ATGCCGGCACAT

### Construction of UAS-DIP transgenic flies

#### Generation of 20xUAS-DIP-γ and 20xUAS-DIP-ε

cDNA of DIP-γ (Flybase Id: FBcl0116341) was ordered from DGRC (No: 7563), and cloned into pJFRC150-20xUAS (Janelia Farm Research Campus) by PCR using the following primers: Forward: ATAGCGGCCGCAAACATGGAGAAGCTGCTGCAATC Reverse: ATATCTAGATCACGCTCCCCCGGTTCTCA The construct pJFRC150-20xUAS-DIP-γ was then inserted into the attp5 genomic site, resulting in the 20xUAS-DIP-γ strain.

cDNA for DIP-ε (Flybase Id: FBcl0317480) was ordered from DGRC (No: 1605414), and cloned into pJFRC150-20xUAS (Janelia Farm Research Campus) by PCR using the following primers: Forward: ATAGCGGCCGCAAACATGGCATACCACCTCGAAGC Reverse: GCATCTAGATCAACCACACGAGTGTGTCG The construct pJFRC150-20xUAS-DIP-ε was then inserted into the attp40 genomic site, resulting in the 20xUAS-DIP-ε strain.

### Antibodies

The primary antibodies used were as follows: anti-V5 (mouse, 1:200) was purchased from Bio-Rad. Anti-DsRed (rabbit, 1:200) was purchased from Clontech Laboratories, Inc. Anti-DIP-beta (guinea pig, 1:300), anti-DIP-epsilon (rabbit, 1:500), and anti-DIP-gamma (guinea pig, 1:400) were gifts from C. Desplan.

The secondary antibodies used were as follows: goat anti-mouse IgG (H+L) Highly Cross-Adsorbed Alexa Fluor 488, Goat anti-rabbit IgG (H+L) Highly Cross-Adsorbed Alexa Fluor 647, Goat anti-guinea pig IgG (H+L) Highly Cross-Adsorbed Alexa Fluor 647 were all purchased from Thermo Fisher Scientific and were all used at a dilution of 1:500.

| Reagent type (species) | Designation | Source or reference | Identifiers | Add. Info. |
| --- | --- | --- | --- | --- |
| Antibody | Anti-V5 (mouse) | Bio-Rad/AbD Serotec | Cat# MCA2892GA; RRID:AB_1658039 | 1:200 |
| Antibody | Anti-DsRed (rabbit) | Clontech Laboratories, Inc. | Cat# 632496; RRID:AB_10013483 | 1:200 |
| Antibody | Anti-DIP-β (guinea pig) | | | 1:300 |
| Antibody | Anti-DIP-ε (rabbit) | | | 1:500 |
| Antibody | Anti-DIP-γ (guinea pig) | | | 1:400 |
| Antibody | Goat anti-Mouse IgG (H&L) Highly Cross-Adsorbed Secondary Antibody, Alexa Fluor 488 | Thermo Fisher Scientific | Cat# A11029 | 1:500 |
| Antibody | Goat anti-Rabbit IgG (H&L) | Thermo Fisher Scientific | Cat# A21245 | 1:500 |
|  | Highly Cross-Adsorbed Secondary Antibody, Alexa Fluor 647 |  |  |  |
| Antibody | 647 Goat anti-Guinea Pig IgG (H&L) Highly Cross-Adsorbed Secondary Antibody, Alexa Fluor 647 | Thermo Fisher Scientific | Cat# A21450 | 1:500 |

### Production of DIP antibodies

DIP-γ antigen: (aa22-393) full length except the predicted signal peptide and the TM domain (guinea pig)GSTQNQHHESSSQLDPDPEFIGFINNVTYPAGREAILACSVRNLGKNKVGWLRASDQTVLALQGRVVT NARISVMHQDMHTWKLKISKLRESDRGCYMCQINTSPMKKQVGCIDVQVPPDIINEESSADLAVQEG EDATLTCKATGNPQPRVTWRREDGEMILIRKPGSRELMKVESYNGSSLRLLRLERRQMGAYLCIASNDVPPAVSKRVSLSVQFAPMVRAPSQLLGTPLGSDVQLECQVEASPSPVSYWLKGARTSNGFASVSTASLESGSPGPEMLLDGPKYGITERRDGYRGVMLLVVRSFSPSDVGTYHCVSTNSLGRAEGTLRLYEIKLHPGASASNDDHLNYIGGLEEAARNAGRSNRTTWQ DIP-β antigen: (23-405aa) full length except a few AAs of the first Ig and the predicted signal peptide and the TM domain (guinea pig) NKISSVGAFEPDFVIPLENVTIAQGRDATFTCVVNNLGGHRVSGDGSSAPAKVAWIKADAKAILAIHEHVITNNDRLSVQHNDYNTWTLNIRGVKMEDAGKYMCQVNTDPMKMQTATLEVVIPPDIINEETSGDMMVPEGGSAKLVCRARGHPKPKITWRREDGREIIARNGSHQKTKAQSVEGEMLTLSKITRSEMGAYMCIASNGVPPTVSKRMKLQVHFHPLVQVPNQLVGAPVLTDVTLICNVEASPKAINYWQRENGEMIIAGDRYALTEKENNMYAIEMILHIKRLQSSDFGGYKCISKNSIGDTEGTIRLYEMERPGKKILRDDDLNEVSKNEVVQKDTRSEDGSRNLNGRLYKDRAPDQHPASGSDQLLGRGTMR DIP-ε antigen: (222-417aa) VDFSPMVWIPHQLVGIPIGFNITLECFIEANPTSLNYWTRENDQMITESSKYKTETIPGHPSYKATMRLTITNVQSSDYGNYKCVAKNPRGDMDGNIKLYMSSPPTTQPPPTTTTLRRTTTTAAEIALDGYINTPLNGNGIGIVGEGPTNSVIASGKSSIKYLSNLNEIDKSKQKLTGS SPKGFDWSKGKSSGSHG Antigens and antibodies were produced at Genescript

### Immunohistochemistry

Fly brains were dissected in Schneider’s medium and fixed in 4% paraformaldehyde in phosphate buffered lysine for 25 min. After fixation, brains were quickly washed with phosphate buffer saline (PBS) with 0.5% Triton-X-100 (PBT) and incubated in PBT for at least 2 hr at room temperature. Next, brains were incubated in blocking buffer (10% NGS, 0.5% Triton-X-100 in PBS) overnight at 4°C. Brains were then incubated in primary antibody (diluted in blocking buffer) at 4°C for at least two nights. Following primary antibody incubation, brains were washed with PBT three times, 1 hr per wash. Next, brains were incubated in secondary antibody (diluted in blocking buffer) at 4°C for at least two nights. Following secondary antibody incubation, brains were washed with PBT two times, followed by one wash in PBS, 1 hr per wash. Finally, brains were mounted in SlowFade Gold antifade reagent (Thermo Fisher Scientific, Waltham, MA).

Confocal imaging was accomplished using either a Leica SP8 laser scanning confocal microscope or a Zeiss LSM800 Laser Scanning Microscope.

### Cell-specific labeling of L2 and L4 neurons in DIP KO experiments (Fig. 1H-O)

(see fly genotype section for exact genotypes)

L2 and L4 split-GAL4 drivers (Tuthill et al., 2013) were used to drive expression of UAS-Flp, which in turn activated the expression of LexA via the Brp BAC used in STaR experiments (see Fig. 2A). In the presence of Flp (i.e. in L2 or L4 neurons) LexA is co-translated with Brp and enters the nucleus turning on the expression of LexAop-myr-tdTOM.

### Quantification of L2 and L4 cell numbers

L2 and L4 cell numbers were determined blind to genotype using cell-specific genetic labeling (described above). Each cartridge in the lamina contains dendrites from a single L2 neuron and the axon/neurite of a single L4 neuron. As cartridges are regularly spaced within the lamina, L2 dendrites and L4 axons within each cartridge can be identified in cross section views of the lamina. The percentage of cartridges containing L2 or L4 neurons was determined for each optic lobe scored. The percentages for lobes of the same genotype were pooled and the average percentage was determined.

### Quantification of Brp puncta in the distal regions of lamina cartridges

Using confocal microscopy, we generated z-stacks of the lamina down the long axis of lamina cartridges. Within each z-stack (i.e. each optic lobe) 25 well labeled cartridges were identified and the number of Brp puncta in their distal halves was counted. The top (distal edge) and bottom (proximal edge) of each cartridge was determined by the first and last sections containing L cell processes (myr-tdTOM), respectively. The midpoint of each cartridge was then identified as the section in between the top and bottom sections. Brp puncta were counted in the sections distal to the midpoint of each cartridge. This stringent criterion was used to avoid counting L2-L4 synapses in the proximal lamina. It is likely that our quantification of distal Brp puncta is an underestimate of the number of ectopic synapses formed in the absence of DIP function. Genotypes were scored in a blind manner by three individuals, and their scores were averaged.

### Quantification of the number of cartridges displaying ectopic synapses in DIP mis-expression experiments

Z-stacks of the lamina were generated down the long axis of cartridges using confocal microscopy. The distal regions of cartridges were determined as in DIP KO experiments described above. Cartridges containing clusters of ≥3 Brp puncta in their distal halves were scored as containing ectopic L cell synapses. 25 cartridges were analyzed per optic lobe, and genotypes were scored in a blind manner.

### Electron microscopy

The heads of 6-day old flies were dissected, immersed in a cacodylate-buffered paraformaldehyde and glutaraldehyde primary fixative, and processed for EM, as previously reported (Meinertzhagen and O’Neil, 1991; Meinertzhagen, 1996). Sections from Epon embedded specimens were cut serially at 60 nm, stained with 4% aqueous uranyl acetate and viewed with a FEI Tecnai 12 electron microscope operated at 80kV, and images collected with a Gatan 832 digital camera. A series of 500 consecutive sections in total was cut, 320 of which were imaged, aligned in Image J, and profiles identified and synapses marked manually.

## SUPPLEMENTAL INFORMATION

**Figure S1.**
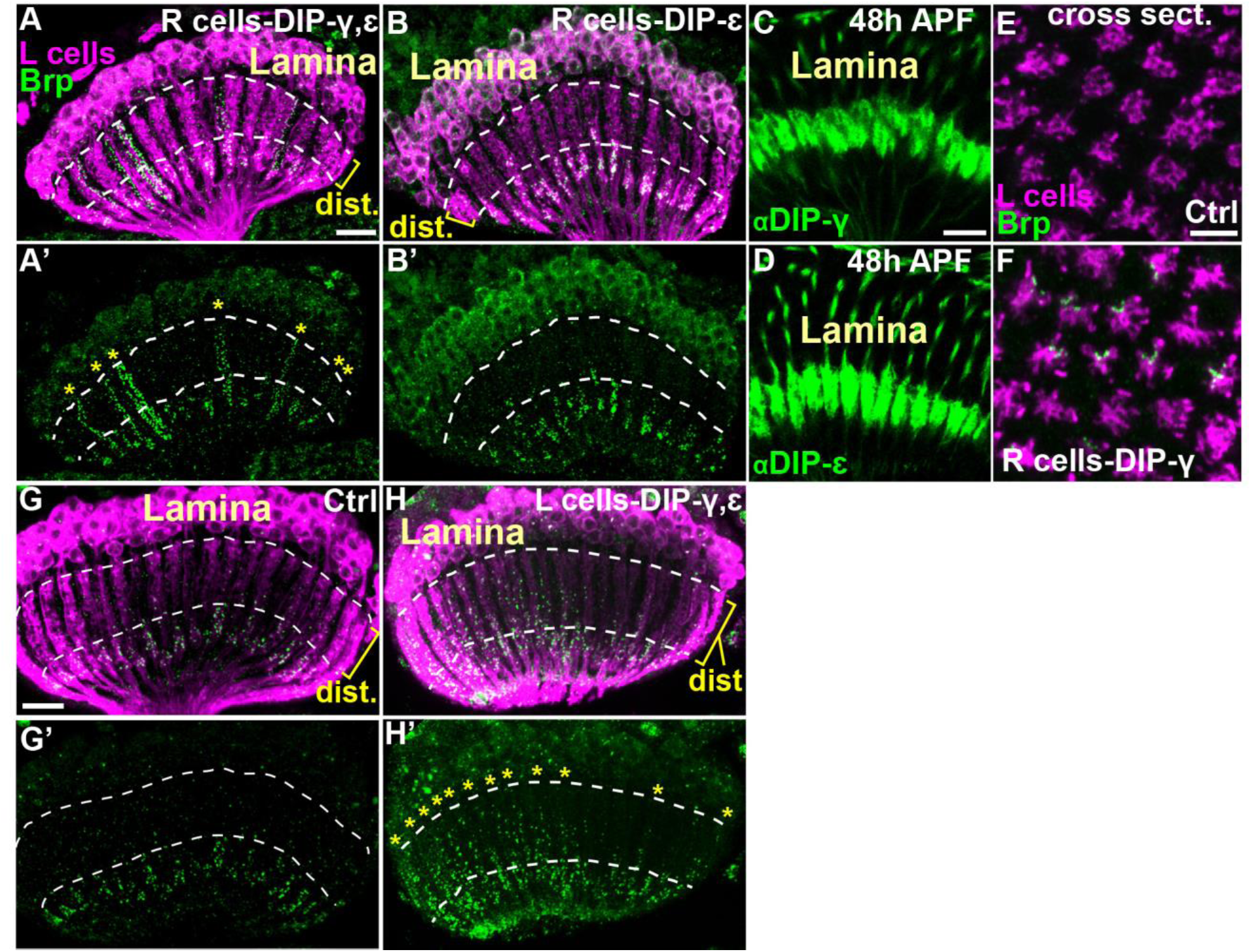
Confocal analysis of DIP mis-expression (related to Figure 4). (A-B’) Confocal images (longitudinal view of lamina cartridges, 1-2 day old adults) show the distribution of Brp (green-smFPV5) expressed in L cells (magenta-myr-tdTOM) in the laminas of flies mis-expressing DIPs-γ and ε or DIP-ε alone in R cells. The area in between the white dotted lines delineates the distal lamina (dist.). Scale bar (A)= 10µm. (A and A’) Mis-expression of DIPs-γ and ε causes L cells to form streams of ectopic synapses within the distal regions of cartridges (yellow stars in A’). n= 9 brains. (B and B’) Mis-expression of DIP-ε only was not sufficient to induce ectopic L cell synapses in the distal lamina. n= 5 brains. (C and D) Immunolabeling of DIPs-γ (green, C) (n=2 brains) and ε (green, D) (n=2 brains) in mis-expression experiments (DIPs-γ and ε in R cells) at 48h APF. Both proteins are strongly expressed in R cell terminals within the lamina neuropil. Scale bar (C)= 10µm. (E and F) Confocal images of cross-sections through the laminas of control flies (GMR-GAL4; n= 3 brains) and flies mis-expressing DIP-γ in R cells (n= 13 brains). The morphology and spacing of L cell processes (magenta-myr-tdTOM) is similar under both conditions. Scale bar (E)= 5µm. (G-H’) Mis-expression of DIPs-γ and ε in L cells (27G05-GAL4). Confocal images show the distribution of Brp (green-smFPV5) expressed in L cells (magenta-myr-tdTOM) in the laminas of control flies (27G05-GAL4 alone) or experimental flies (27G05-GAL4 and UAS-DIP-γ and ε). The region between the dotted lines delineates the distal lamina (dist.). Scale bar (G)= 10µm. (G and G’) Brp is restricted to the proximal lamina (beneath the distal lamina) in control flies. n=5 brains. (H and H’) L cells form ectopic synapses in the distal regions of lamina cartridges (yellow stars in H’) upon mis-expression of DIPs-γ and ε in L cells. n= 5 brains.

**Figure S2.**
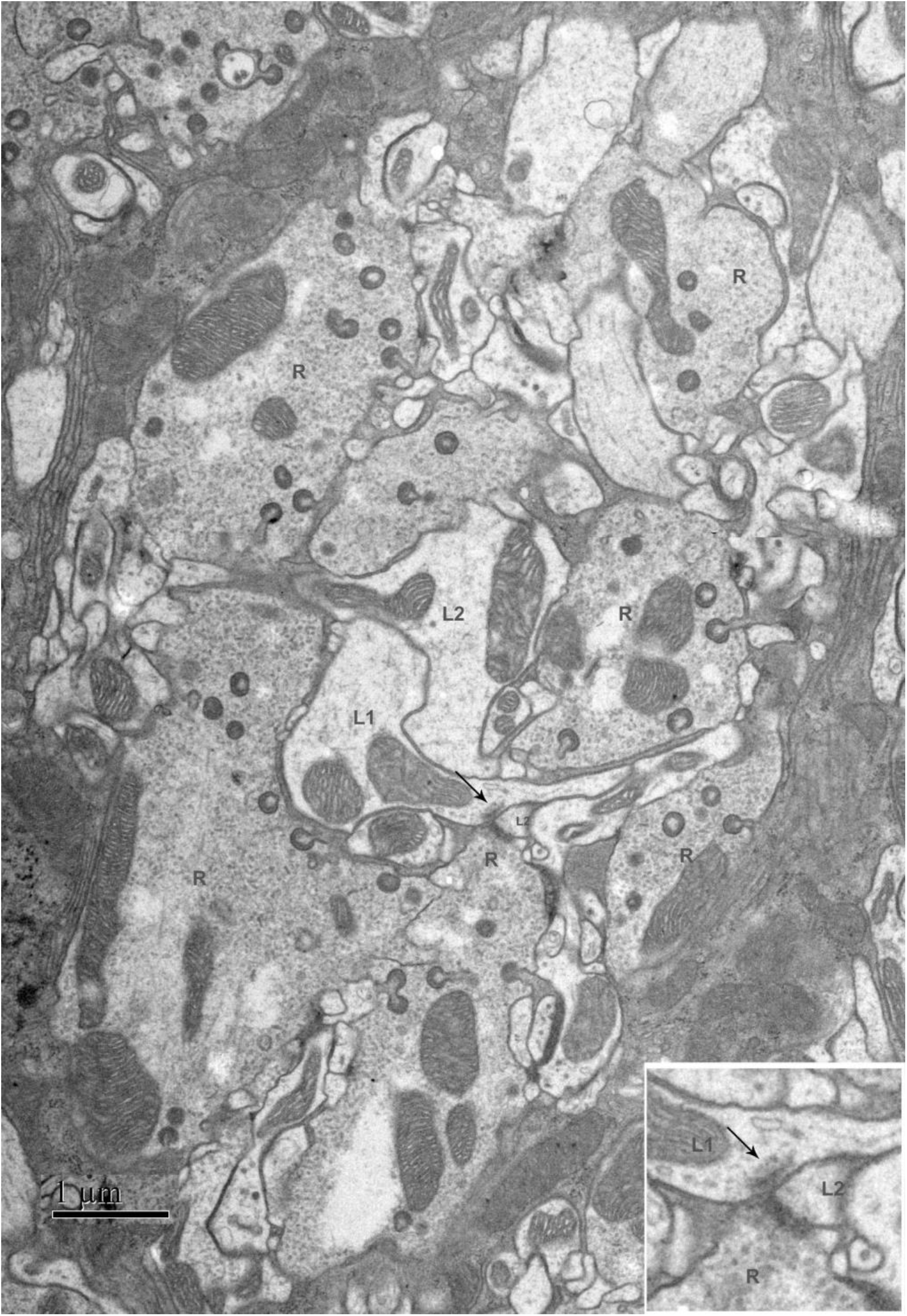
EM analysis of DIP mis-expression (related to Figure 4). Cross section through a lamina cartridge (60nm section) from a fly mis-expressing DIPs-γ and ε in R cells imaged by TEM. 6 R cell profiles surrounding the axon profiles of L1 and L2 neurons are identified, demonstrating that the general arrangement of profiles within cartridges is not perturbed under these conditions, allowing each profile to be identified from its position. Arrow at pre-synaptic site indicates a putative L1 to R cell synapse (also shown in Figure 4E). Scale bar= 1µm.

**Table S1. EM analysis of DIP mis-expression Related to Figure 4.** Numbers and distributions of R and L cell pre-synaptic sites identified by EM within a cartridge from a fly mis-expressing DIPs-γ and ε in R cells. The first row indicates pre-synaptic cell types. The left most column (Layer) shows the positions of synapses within the cartridge from distal (Layer 1) to proximal (Layer 311). Each layer reports data for x10 60nm sections, Layer 1 thus comprising sections 1-10. If processes could not be traced throughout the cartridge B indicates the section where it was first identified, and E indicates the last section in which it was identified.

